# New realm of precision multiplexing enabled by massively-parallel single molecule UltraPCR

**DOI:** 10.1101/2023.10.09.561546

**Authors:** Janice H. Lai, Jung won Keum, Haeun G. Lee, Mehdi Molaei, Emily J. Blair, Sixing Li, Jesse W. Soliman, Vedant K. Raol, Camille L. Barker, Stephen P.A. Fodor, H. Christina Fan, Eleen Y. Shum

## Abstract

PCR has been a reliable and inexpensive method for nucleic acid detection in the past several decades. In particular, multiplex PCR is a powerful tool to analyze many biomarkers in the same reaction, thus maximizing detection sensitivity and reducing sample usage. However, balancing the amplification kinetics between amplicons and distinguishing them can be challenging, diminishing the broad adoption of high order multiplex PCR panels. Here, we present a new paradigm in PCR amplification and multiplexed detection using UltraPCR. UltraPCR utilizes a simple centrifugation workflow to split a PCR reaction into ∼34 million partitions, forming an optically clear pellet of spatially separated reaction compartments in a PCR tube. After *in situ* thermocycling, light sheet scanning is used to produce a 3D reconstruction of the fluorescent positive compartments within the pellet. At typical sample DNA concentrations, the magnitude of partitions offered by UltraPCR dictate that the vast majority of target molecules occupy a compartment uniquely. This single molecule realm allows for isolated amplification events, thereby eliminating competition between different targets and generating unambiguous optical signals for detection. Using a 4-color optical setup, we demonstrate that we can incorporate 10 different fluorescent dyes in the same UltraPCR reaction. We further push multiplexing to an unprecedented level by combinatorial labeling with fluorescent dyes — referred to as “comboplex” technology. Using the same 4-color optical setup, we developed a 22-target comboplex panel that can detect all targets simultaneously at high precision. Collectively, UltraPCR has the potential to push PCR applications beyond what is currently available, enabling a new class of precision genomics assays.

## INTRODUCTION

Since its invention, PCR has become an indispensable tool in modern biology (1–9). It is fast, inexpensive, has high-specificity, and has become a standard for quantifying DNA targets. Because of these qualities, PCR has become the *de facto* gold standard for validating the results of other technologies and for detecting DNA in clinical medicine.

A particular challenge in PCR-based detection assays is how to multiplex for a higher number of DNA targets (6, 10–13). In most cases, a fluorescent dye is used to label either double-stranded DNA or a specific PCR product when conjugated to an oligonucleotide probe (14–17). This simple approach where one color is assigned to one target generates robust, easy-to-detect fluorescent signals and is almost universally used (6, 13). Widely adopted yet fluorescence based PCR target detection is largely restricted to small numbers of different molecules (typically less than 4) per reaction. Indeed, although there is a large repertoire of useful fluorescent labels (18), simply increasing the number of primers and colors as an attempt to detect multiple targets is fraught with complexity due to inter-molecular competition for reagents and primers and results in a complex and intractable overlay of spectral overlaps (6, 13–16).

We have previously demonstrated that UltraPCR has an outstanding counting capability with dynamic range and single molecule precision to rival that of next generation sequencing (NGS) (19). In this study, we show how UltraPCR’s singlet amplification leads to clean signal amplification and detection (**Figure 1A**), which enables high order multiplexing and a straightforward data analysis workflow. First, UltraPCR’s emulsion and imaging systems unlock the ability to incorporate 10 fluorescent dyes into a single PCR assay. Second, singlets enable combinatorial labeling without the statistical limitations of multiple target occupation seen in typical dPCR technologies (19–22) (**Figure 1B**). Third, the quasi-solid phase optically clear emulsion pellet allows for repeated interrogation via 3D imaging of the large number of partitions. When combined, these features allow for a highly scalable, inexpensive approach to high-level PCR multiplexing. Accordingly, UltraPCR catapults multiplex PCR capabilities to an unprecedented level of complexity while achieving extremely high precision. We demonstrate that the combination of these innovations can build high-plex, high-precision assays by showcasing an 8-color, 22-plex pathogen ID panel as a proof-of-principle application.

**Figure 1.**
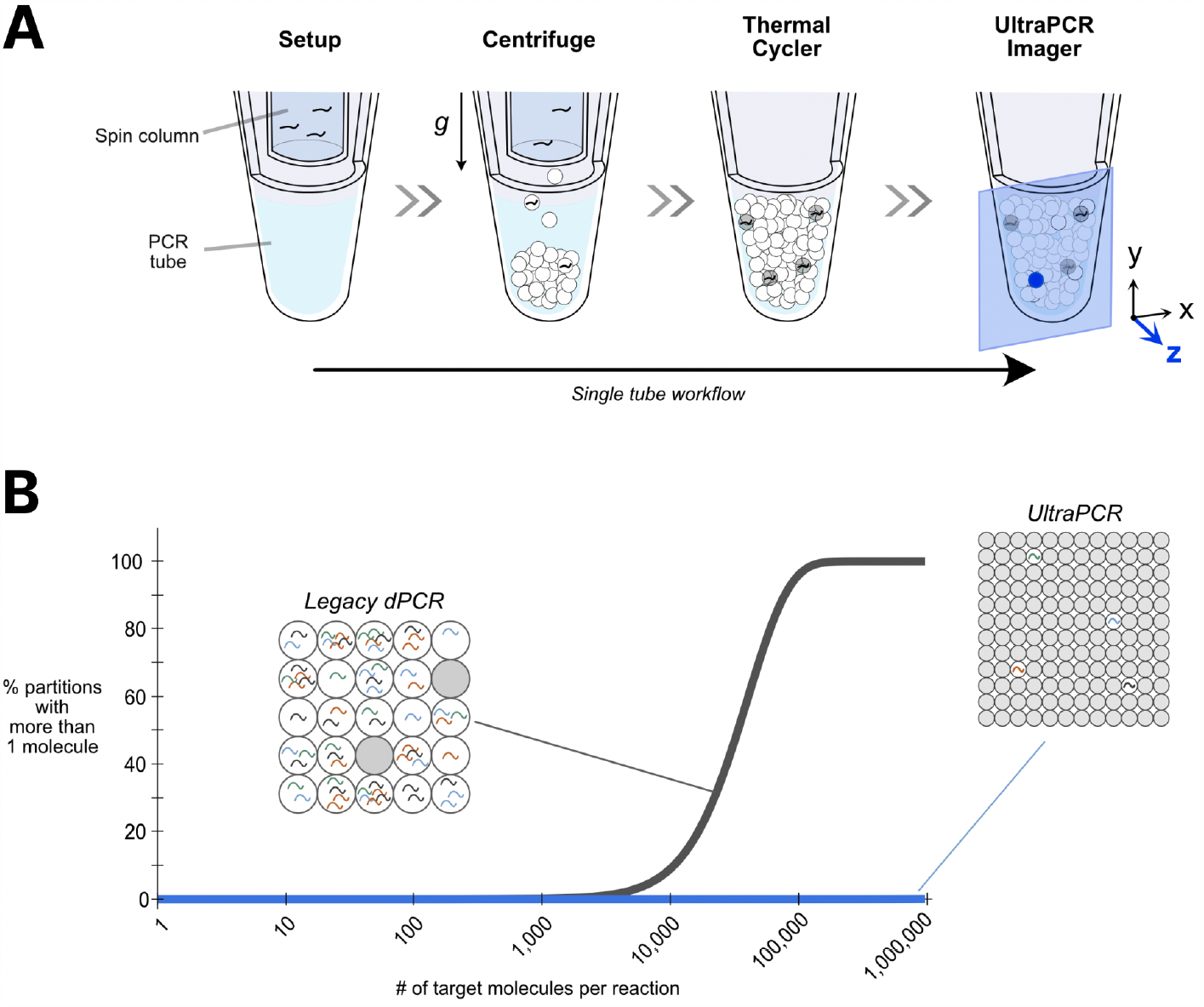
Overview of UltraPCR. (A) Workflow diagram outlining how UltraPCR works in a 4-step process. Step 1: PCR mix is loaded onto the spin column. Step 2: Centrifugation draws the PCR mix through a custom-designed column to divide a 50 uL reaction into ∼34 million partitions within 20 minutes. Step 3: Thermal cycling of the UltraPCR emulsion amplifies signal in partitions with single molecule templates. Step 4: UltraPCR Imager analyzes the optically clear UltraPCR emulsion by translating a laser light sheet through the PCR tube. (B) Plot displaying the % of partitions encompassing more than 1 molecule in legacy dPCR systems (20,000 partitions, black) (20, 25, 26) vs UltraPCR (34,000,000 partitions, blue) (19). In UltraPCR, even at 1,000,000 molecule input, ∼99.95% of partitions are either empty or with single target molecules, which enables a paradigm shift of massively parallel single molecule PCR. Schematic of UltraPCR DNA occupancy mimics the actual proportion of partitions with 1 million molecules in 34 million partitions (2.9% occupancy). On the other hand, in legacy dPCR systems, even with 50,000 molecule input, more than 70% of the partitions are occupied with two or more target molecules (250% occupancy), as shown in the schematic.

## RESULTS

### Optical signature profiling unlocks additional fluorescent dye repertoire for PCR

UltraPCR provides a fundamentally different approach to fluorescent dye labeling, readout, and identification for multiplexed assays. During centrifugation-based partitioning (23), each DNA target molecule of a sample is randomly partitioned into individual singlets directly into the PCR tube (**Figure 1A**). Collectively, UltraPCR’s unique chemistry renders emulsions optically clear with spatially immobilized droplets/partitions, facilitating the use of 3D light sheet microscopy to measure fluorescence signals of individual partitions in a massively parallel manner with high spatial resolution (19) (**Figure 1A**). Currently, a single light sheet channel takes ∼40 seconds to sweep through a sample of ∼34 million partitions. The static state of emulsion enables repeated imaging where the 3D spatial location of partitions remains traceable without sample manipulation. Leveraging these features, UltraPCR can perform serial imaging of the same PCR tube with different “channels” — each channel having its own excitation and emission configuration — to generate an “optical signature” of each positive partition (**Figure 2A**).

**Figure 2.**
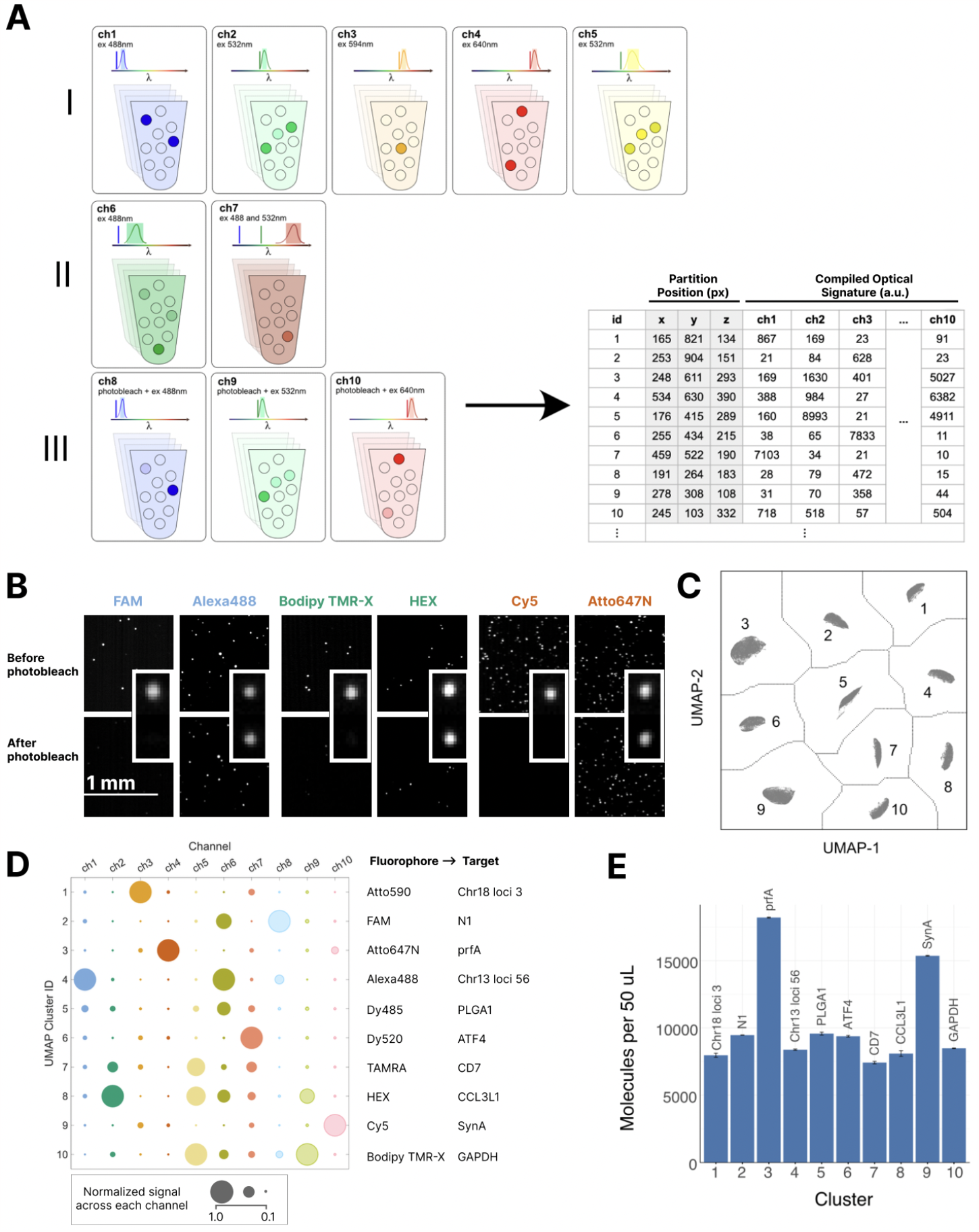
Generation of optical signature with spatial information using 10 dyes. (A) Schematic of how optical signature is captured. In this specific example, every partition is scanned serially with 10 different channel settings configured for dye classes I, II, and III. Channel settings include distinct excitation wavelength, indicated by colored vertical lines, emission wavelength, indicated by colored rectangles, and photobleaching events. Data generated in each channel is used to assemble an optical signature for each partition composed of fluorescence intensity values, which unambiguously identifies the target captured in 3D partition positions. Sample of data is shown in the table (B) light sheet images with photobleachable (FAM, Bodipy TMR-X, and Cy5) and photostable (Alexa488, HEX, and Atto647N) dyes before and after a brief photobleaching event. Zoomed in micrographs of individual partitions are shown in the corner of images as examples. (C) UMAP visualization of optical signatures for all positive partitions detected by the UltraPCR Imager analysis pipeline followed by clustering using DBScan (27). (D) Aggregate optical signature for each cluster (vertical-axis) with circle size indicating normalized intensity value for each channel (horizontal-axis). (E) Number of molecules identified per cluster. This experiment was performed as a duplicate to generate an error bar signifying standard error.

The concept of an optical signature enables the profiling of highly similar fluorescent dyes that may otherwise be difficult to distinguish in traditional PCR platforms. For example, HEX and TAMRA dyes are typically not used concurrently in qPCR/dPCR due to their high spectral overlap (24). However with UltraPCR, these dyes can be differentiated in 2 serial imaging steps. Using the same excitation laser (532 nm), we created 2 channels in UltraPCR that employ different emission bandpasses for the detection of HEX (ch2) and TAMRA (ch5) (**Figure 2A**). Even though the emission of HEX and TAMRA dyes are detectable in both channels, their fluorescence intensities in each channel collectively render distinct optical signatures.

We set out to identify more dyes that the UltraPCR Imager can uniquely measure. We were able to distinguish three major dye classes for a total of 10 dyes (that are commercially available at the time of publication) that are compatible with the UltraPCR Imager configuration (**Figure 2A**), using different combinations of excitation lasers, emission filters, and imaging settings (also referred to as “channels”). The first class of dyes include Alexa488, HEX, Atto590, Atto647N, and TAMRA (ch1, ch2, ch3, ch4, ch5, respectively).

The class II dyes (Dy485XL (ch6) and Dy520XL(ch7)) use the same excitation lasers as the class I, but because they have larger Stokes shifts (28, 29), the optical signatures can be uniquely identified by creating channels with the same excitation laser but different emission filters. The class III dyes are differentiable based on their photobleaching properties (30, 31). We identified 3 additional dyes that have similar optical signatures as class I dyes, but can be photobleached in a controllable manner. Light sheet images of positive partitions labeled with these two classes of dyes reveal their distinctive optical signature (**Figure 2B**); these class I and III dye pairs are FAM (photobleach sensitive, ch8) vs Alexa488, Bodipy TMR-X (photobleach sensitive, ch9) vs HEX, and Cy5 (photobleach sensitive, ch10) vs Atto647N. We added a photobleach event using UltraPCR Imager light sheet between 2 imaging events of the same excitation/emission setting that allows us to easily differentiate these dye pairs (**Figure S1A**).

As a proof-of-principle, we developed a 10-plex UltraPCR panel using our new dye differentiating approach and these 10 dyes. Positive partitions for each channel were automatically detected by signal intensity difference from background using a proprietary algorithm supplied by the UltraPCR Imager. Optical signatures from 10 channels were gathered to generate a vector of intensity values [ch1-ch10] for each positive partition detected in different locations [x,y,z] of the tube (**Figure 2A**). This 2D matrix was then used to visualize partitions using Uniform Manifold Approximation and Projection, UMAP (32), followed by clustering to distinguish the targets (**Figure 2C**). Using these techniques, we could quickly distinguish 10 clusters of positive partitions with similar optical signatures in an unsupervised manner. The clear separation of clusters in UMAP showcases the difference in optical pattern of partitions with different dyes. We examined the optical signature of each cluster by measuring median signals of partitions residing in them to identify the fluorophore (and thereby the amplicon target) represented by the cluster (**Figure 2D**). The data shown for each channel is normalized by the maximum signal across each channel. Since each datapoint was represented by a single target molecule, the molecule count per target was simply the number of datapoints per cluster (**Figure 2E**). The molecule counts per cluster in the 10-plex assay matched the expected counts based on a singleplex assay with no observable differences in counting precision (**Figure S1B**). We were able to demonstrate that singlet partitions paired with optical signature profiling can increase multiplex capacity in PCR.

### Comboplex: combinatorial labeling of singlet PCR dramatically expands multiplexing capacity

The multiplex capacity can be extended when we combine dyes combinatorially, in a method we call “comboplex,” creating new distinctive optical signatures. As a proof-of-principle, we labeled a single target with either 1, 2, or 3 fluorescent labels where each configuration targets the same gene using the same probe sequence but conjugated to different dyes (**Figure 3A**). Using a set of 4 dyes, we tested all 14 combinations where each comboplex assay was serially imaged in 4 channels (ch1-4) to collect fluorescent intensities (**Figure 3A**). In this particular configuration of the assay, even though the fluorescence intensity of targets with more than one fluorophore decreased due to probe competition for the same target amplicon, we were still able to call positive partitions apart from background (**Figure S2A**). We first analyzed positive partitions in each channel independently and saw that the molecule counts for all label combinations were highly comparable (**Figure S2B**). Another way of analyzing comboplex data was to combine the fluorescence optical signatures into a multivariable matrix and use UMAP and automatic clustering to identify dye combinations (**Figure 3B**). The separation of the clusters with combinatorial labels is comparable with the separation of clusters with single dye per target (**Figure 2C**) indicating efficacy of comboplex assays. Each of the 14 clusters identified in the UMAP also have unambiguous optical signatures that can be used to map the actual dye combination used (**Figure 3C**).

**Figure 3.**
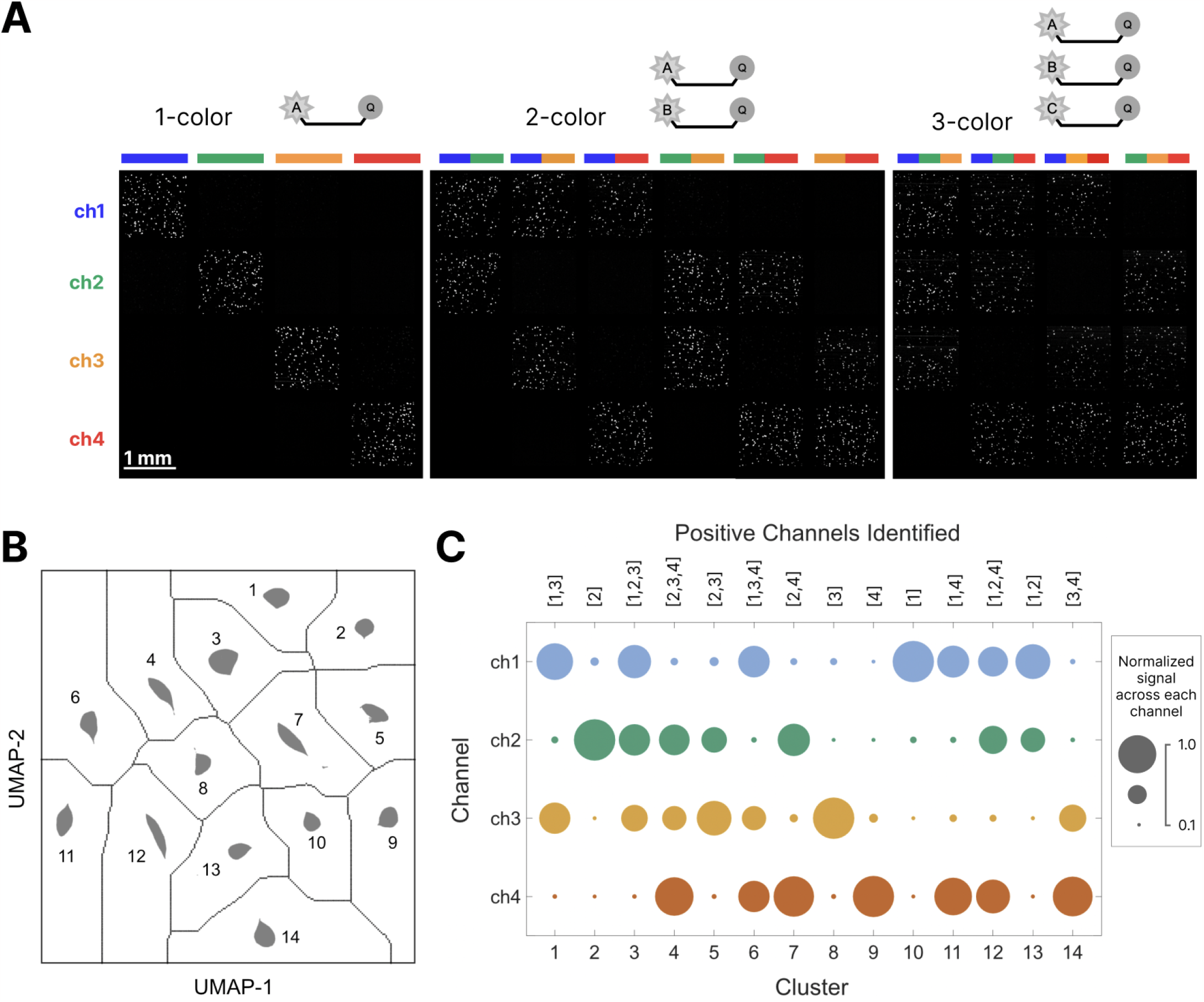
Comboplex overview. Strategy for labeling a target with more than one dye. (A) In this design, a target was amplified using 1 to 3 hydrolysis probes using the same sequence but conjugated with different color dyes (shown in the bar above the image, labeled A, B, and C), amplifying a gene segment of prfA of L.cytomonogenes, resulting in 1 color, 2 color, or 3 color combinations. Sample light sheet images in each channel shown in different rows display positive partitions in each color combination. (B) Each color combination generates a unique optical signature that can be visualized on UMAP. (C) Plot of each channel’s fluorescence intensity, indicated by size of the circles, for each comboplex configuration; cluster numbers in the horizontal axis correspond to cluster numbers on UMAP.

The concept of comboplex complements our singlet optical signature technique, as the same set of “base” fluorescent labels can be used to massively expand the configurations for single molecule labeling. The number of comboplex targets can be calculated by the following equation: *P* (*n, k*) = *n*! */* (*n*−*k*)!, where *n* is the number of fluorophores available for the platform, and *k* is the number of labels used per target. Combined with the possibility that a target can be labeled with 1 or 2 fluorophores (*k* = 1 or 2), the upper limit of comboplex can be extremely high. This approach is particularly useful because UltraPCR has at least a 10 dye compatibility as demonstrated above. Therefore, the theoretical plex can reach >50 if 1 and 2 dye combinations are used, which is at least 10-fold higher than typical PCR, as outlined in **Table 1**.

**Table 1.** Possible number of multiplex combinations achievable by the comboplex approach.

| # of dyes (n) | k=1 | k=2 | k=3 | k=1, 2 or 3 |
| --- | --- | --- | --- | --- |
| 4 | 4 | 6 | 4 | 14 |
| 5 | 5 | 10 | 10 | 25 |
| 6 | 6 | 15 | 20 | 41 |
| 7 | 7 | 21 | 35 | 63 |
| 8 | 8 | 28 | 56 | 92 |
| 9 | 9 | 36 | 84 | 129 |
| 10 | 10 | 45 | 120 | 175 |

### Demonstration of simultaneous digital counting of 22 targets in a single tube

We leveraged our expanded dye repertoire to build a proof-of-concept comboplex panel in UltraPCR. We selected a set of 22 common targets for identifying respiratory pathogens and associated antibiotic resistance genes (33). In our multiplex design, we used the top 8 most commercially available DNA dyes to label each target with either 1 or 2 unique labels using a Universal Multiplex strategy (**Figure 4A**). These multiplex primers were designed *in silico* using a custom primer design algorithm with the principles to minimize primer dimers and non-specific primer extensions to maximize compatibility. Each target labeling strategy was confirmed in single target reactions before pooling together for multiplexing.

**Figure 4.**
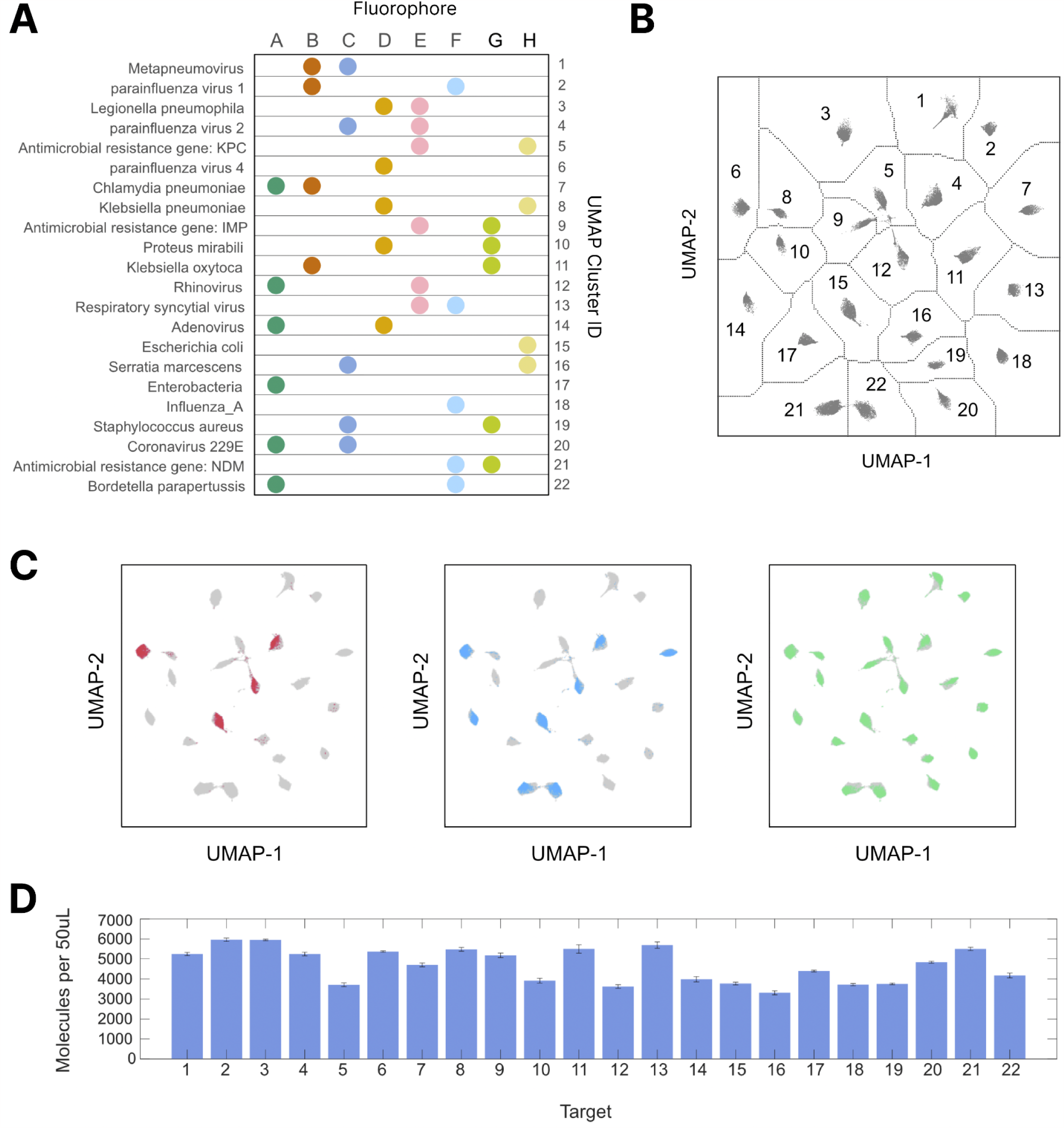
Ultra-high multiplex panel using comboplex in UltraPCR. (A) Design of a 22-plex pathogen identification panel, showing the assignment of the 8 fluorescent dyes (labeled A, B, C, D, E, F, G, H, each associated with a unique dye). (B) Simultaneous visualization of the reference optical signatures of each of the 22 targets using UMAP. This reference map was generated by combining optical signatures of all positive partitions from 22 UltraPCR reactions, each performed with one of the 22 synthetic template targets. (C) Different numbers of targets (left) 4, (middle) 8, (right) 22 were spiked into the reaction to show that the reference signatures can be used for pathogen identification without the need for manual analysis and target assignment as in conventional dPCR. Each partition in the sample with all 22 targets was mapped to the nearest clusters of the reference signature to derive the molecule count detected per target in (D). Bar graph displaying the average molecule counts for each of the 22 targets all present in the reaction across 4 technical replicates, and the error bar displaying the standard error.

With this magnitude of multiplex capability, we designed a new analysis workflow to streamline data interpretation. We generated a ‘reference optical signature’ for each of the 22 targets by performing the 22-plex UltraPCR assay on spike-in DNA of each target individually. Each of these reactions was imaged through 8 channels to record the collective optical signature for each target. All optical signatures were combined and the 8-dimensional data was projected onto a 2D graph using UMAP. Twenty-two distinct clusters were identified using an automatic clustering algorithm (**Figure 4B**). Using this automated approach, we avoided tedious manual determination of target assignment for each partition, a method still commonly applied in legacy systems (34). Note that this map of the 22-plex assay can be applied across multiple experiment days and probe batches, further streamlining the analysis of these higher order multiplexing samples.

We subsequently used this ‘reference map’ to determine the identities and molecule counts of targets in samples. In the first experiment, we added 2 targets concurrently at different concentrations; target 1, *Imp*, was spiked at a constant level whereas target 2, *Adv54*, was serially diluted. Each reaction was analyzed independently using our “cluster-counting” workflow aforementioned. We observed that even at wide variation in spike-in levels of these 2 targets, the precision of molecule counting by UltraPCR remains extremely high (**Figure S3**). The data is consistent with the notion that even at such a high order of “multiplexing”, UltraPCR remains functional at a singlet realm where amplification and detection remain independent between different target molecules.

We extended the singlet multiplexing notion further by spiking in more targets concurrently into a 22-plex comboplex assay. We tested replicates where 4, 8, or all 22 target templates were added into the assay; the optical signature of each positive partition was overlaid with the reference map to show little batch-to-batch variability in cluster identification, even though this spike-in experiment was performed on a separate day using a separate batch of fluorescent probes for some targets. In all conditions, we were able to simultaneously detect the presence of multiple targets and quantify these targets (**Figure 4C**). Additionally, for the 22-target test, our 4 technical replicates showed a remarkably high counting precision (**Figure 4D**).

## DISCUSSION

UltraPCR is a novel platform that transforms how DNA can be quantified. We introduced UltraPCR which utilizes millions of picoliter partitions per sample as a next generation PCR technology (19). Each sample is divided into ∼34 million picoliter-sized partitions using a simple centrifugation step (**Figure 1A**) — a 1500-fold higher number of partitions than legacy dPCR (20). At this magnitude, the ratio of DNA to partition is so small that Poisson statistics (35–37) indicate that target molecules are individually cast into separate partitions, giving rise to single molecule partitions — or singlets — in a massively parallel manner (**Figure 1B**). In this realm, PCR within each partition amplifies a single template molecule without competition. Fundamentally, UltraPCR is similar to single molecule PCR demonstrated via limiting dilution (7), but in a massively parallel manner such that even up to 1 million molecules can be housed individually and counted without the use of Poisson correction (**Figure 1B**). The precision and dynamic range of UltraPCR is so far beyond the capability of legacy dPCR (**Figure 1B**) that challenging assays such as fetal aneuploidy detection become possible with as low as 4% trisomy DNA against the backdrop of a normal individual’s background DNA. Our previous study showcased, for the first time, a PCR platform achieving NGS-like counting precision (19). The advantages of singlet partitioning, however, extends beyond counting capabilities; in this study, we show that singlet partitioning itself simplifies the biochemistry of multiplex PCR, enabling a new era towards straightforward and higher-order multiplexing.

Although the need to expand multiplex capabilities has been recognized since the invention of qPCR, a primary limitation is the number of dyes that can be used concurrently per assay (6). In both bulk qPCR and dPCR regimes, partitions inevitably contain mixed targets due to limited partition number (n=1 for qPCR and n∼20,000 for dPCR) (**Figure 1B**). Each dye assigned per target therefore must be distinct by design with minimal spectral overlap (6). Due to this limitation, typical dPCR platforms can only support 2-6 colors (20, 34, 38). Even at this low order of multiplexing, emission crosstalk between fluorophores is an intrinsic issue requiring various compensation techniques to remove false positive fluorescence signals. In addition to a low number of concurrently usable fluorophores for multiplex PCR, the amplification signal must be extremely high such that the biomarker signal rises above what is removed during compensation (6).

The UltraPCR platform reimagines how dyes can be used for multiplexing by leveraging our method of optically profiling singlet partitions. Our optical signature data is comparable to gene expression analysis in single cells — where each partition can have fluorescence expression across many imaging channels — and can be collectively analyzed like NGS datasets (**Figure 2**). We applied our proof-of-concept datasets to multivariable analytic tools such as UMAP and associated clustering techniques (32), showcasing how future PCR analysis of many biomarkers can be just as easily automated, with minimal manual intervention for high throughput, high-plex assays (**Figure 2-4**).

This study marks the beginning of expanding the number of dyes that can be used concurrently in a PCR reaction. Just in this pilot study alone, we were able to demonstrate the simultaneous use of 10 commercially available dyes that can be conjugated to DNA probes (**Figure 2**). We believe that the repertoire of dyes that can be used in UltraPCR can be further expanded by at least 2 ways: 1) increasing the number of lasers from 4 to a higher number, matching that of high-end flow cytometers or 2) expanding the number of off-the-shelf dyes that can be conjugated to DNA probes. Consider the number of lasers and dyes observed in flow cytometry: as many as 9 lasers and 50 targets (or parameters) have been commercially achieved (39). The compatibility of these lasers and dyes would need to be explored in the future as more dyes are conjugated to DNA probes for their compatibility to UltraPCR.

Beyond fluorescent dye availability, the performance of traditional multiplex PCR is often complicated and unpredictable as simultaneous amplification of multiple targets can be complex. While the variables of maximizing amplification efficiency — such as primer and probe melting temperatures, the selection of high fidelity enzymes, etc. — are generally understood, PCR efficiency between multiplex targets can be different, leading to ambiguous signals (6). In qPCR and dPCR, suboptimal multiplex assays can manifest in fluorescence signal reduction or Ct value increase, both of which can lower assay sensitivity, accuracy, and reproducibility (13, 25). Extensive work would need to be performed on balancing the amplification success of different targets concurrently and require different parameters for different applications (6, 40, 41). The amplification competition between multiplex targets is not impossible to solve, but can take significant development and optimization to achieve acceptable results. As such, multiplex PCR assays are often only deployed in high throughput settings, like clinical diagnostics, or packaged in verified panels to offset the high cost of assay optimization efforts (13, 33, 38, 42). Users in other applications are left to run concurrent 1-plex PCR reactions to lower costs and assay development overhead or to perform complicated and costly workflows such as NGS.

Several groups have taken on the challenge to increase multiplex capacity in PCR beyond the number of dyes, but with drawbacks (22, 25, 43–45). The first general approach is amplitude-based multiplexing, in which at least 2 targets are detected using the same channel while controlling target-specific amplitude of partitions (25, 43). In some cases, up to 10 targets can be added within the same color space (44). While this method appears attractive on paper, the complexity of multiplexing in these platforms can be complicated during assay optimization and data analysis. First, even more rigorous assay development is required so that different targets can be easily distinguishable within the same optical channel. Second, key reagents for these assays require meticulous quality control measures to prevent issues arising from batch variability effects for each dye probe involved. Data analysis of these samples is further computationally taxing, and requires additional steps for deconvolution of ambiguous partition clusters. Consider a typical dPCR system (∼20,000 partitions per reaction) where partitions are overloaded with 0, 1, or >1 target molecules (**Figure 1B**). In this regime, the number of target molecules cannot be directly counted; rather, statistical techniques that leverage Poisson statistics are heavily used to derive the estimated concentration of DNA targets (20, 22, 46). When assays are multiplexed via amplitude difference where each color harbors 2 targets, at least 3 different intensity thresholds must be set per color to separate partitions that are negative, target 1 only, target 2 only, and target 1 and 2 together (25). In these dPCR systems, the number of thresholds and potential partition clusters increases exponentially with the number of targets (n), where the number of clusters is 2^n^. For example, the theoretical cluster number that needs to be designed in a 12-plex, 6-color dPCR assay is 4096, making it extremely difficult for any user to decipher. Altogether, the amplitude based multiplexing approach is challenging to implement and its application remains niche.

A second multiplex approach demonstrated recently by a dPCR group showcased a concept called color coding, which achieves 15-plex using 6 dyes (20, 22, 26, 46). Color coding in dPCR is subjected to the limitations of multiplet partitions. As eluded by the authors, as the proportion of multiplets increases, mixtures of targets with color codes can generate “ambiguous clusters” that need to be discarded from Poisson correction (20, 22, 26, 46). The presence of these ambiguous clusters fundamentally limits the effectiveness of color coding given that the partitioning capacity of these dPCR systems is already low.

We demonstrated in this study how UltraPCR’s singlet amplification overcomes the many challenges of PCR multiplexing and pushes multiplexing to an unprecedented level. We showcased, by having an expanded dye portfolio and combinatorial labeling enabled by singlet realm, the development of a 22-plex pathogen ID panel for the detection of common respiratory viruses and antibiotic resistance genes (**Figure 4**) (33). In theory, the maximum multiplex capacity is much higher than 22; if we utilize all 10 dyes, where each target is labeled with either 1 or 2 dyes, the maximum limit would be 55-plex (**Table 1**).

The unprecedented level of multiplexing and simultaneous detection of targets can be applied to many applications, ranging from basic research, translational research, to clinical diagnostics. UltraPCR’s multiplexing capability can be useful in gene expression studies and biomarker discoveries, where panels of biomarkers identified by microarray or RNA-seq studies can be quickly and simultaneously validated with a PCR based readout with high precision, reducing the amount of sample usage and number of reactions required. In infectious disease diagnostics, the ability to accurately quantitate one or more (in co-infection cases) pathogens with UltraPCR allows clinicians to determine which pathogen(s) are present, providing potentially actionable results for quick, targeted treatment selection (47, 48). In comparison, traditional microbiology culture techniques take several days longer in turnaround time with no quantitative result on pathogen level (47). In oncology, where the ever-increasing treatment options require larger diagnostic biomarker panels to guide treatment decisions (49, 50), we anticipate that this level of multiplexing capacity offered by UltraPCR can be used to displace certain amplicon sequencing panels, providing broader patient access by dramatically reducing turnaround time and cost of each test as compared to NGS. The additional workflow advantages of UltraPCR — zero dead volume and high PCR loading volume (>30 uL) (19) — would be particularly beneficial in liquid biopsy and rare molecule detection applications.

In conclusion, we provide a foundational set of data showcasing the capability of UltraPCR in both multiplexing and precision counting. Not only can UltraPCR perform typical qPCR/dPCR applications, but it also goes beyond established limitations and fundamentally shifts PCR dynamics due to the singlet realm that UltraPCR uniquely operates within. In UltraPCR, multiplexed assay design and data analysis are dramatically simplified as each DNA molecule is partitioned and amplified separately without the competition of other DNA targets. Altogether, massively parallel single molecule amplification by UltraPCR enables a new class of genomics that is rapid, precise, and inexpensive for translational research and beyond.

## MATERIALS AND METHODS

### UltraPCR workflow

All UltraPCR reaction mixes were prepared using 4X UltraPCR mix (Enumerix) according to the manufacturer’s protocol. For every sample, 50 μL of the UltraPCR reaction mix (with primers, probes, and DNA template) was added to the UltraPCR Spin Columns (Enumerix), outfitted into PCR strip tubes carrying emulsifying reagents (Enumerix). The strip assembly was loaded into a custom UltraPCR swing bucket (Enumerix) for use in an Eppendorf 5430 centrifuge. Up to 48 samples were spun for 20 min at 16,000g to form UltraPCR emulsions. After centrifugation, the spin columns were discarded, and the PCR tubes containing the emulsions were sealed and placed into a Bio-Rad C1000 thermal cycler. The same PCR strip tubes were then placed into an UltraPCR Imager (Enumerix) for positive partition scanning, where a laser light sheet was translated across the PCR tube and the illuminated partitions were imaged. Four lasers were utilized to scan up to 10 dyes, where each dye had an optimized imaging setting with defined excitation laser(s) and emission filters. The current configuration of the UltraPCR Imager includes excitation wavelengths at 488 nm, 532 nm, 594 nm, and 640 nm.

For each sample, cross sectional images of the PCR tube containing emulsions collected by light sheet imaging were compiled to generate a 3D reconstruction of the PCR tube to count positive partitions using custom software written in C# and MATLAB. Given that signal from UltraPCR represents single molecules, the UltraPCR Imager analysis pipeline is deployed without user intervention for positive partition identification. In all samples, an auto-thresholding algorithm was applied to gate signal that is DNA-positive versus negative. Multivariable analysis was performed in MATLAB, using a UMAP package for visualization, and DBScan (27) for clustering.

### Samples

Synthetic templates were obtained as gBlocks ordered from Integrated DNA Technologies. All serial dilutions and template dilutions used TE buffer (Teknova, 10 mM Tris-HCL, 0.1 mM EDTA, pH 8.0) to reach intended copy numbers and confirmed using UltraPCR prior to multiplex experiments.

### Assay designs

Primers and probes were purchased from Integrated DNA Technologies except for PLGA1-Dy485XL and ATF4-Dy520XL probes that were obtained from Biosynthesis. Unless specified, individual primers and probes were designed using Primer3 (51–53). For multiplex assays, primer candidates designed by Primer3 were screened and selected for the multiplex panel using a custom algorithm with the principles to minimize primer dimers and non-specific primer extensions to maximize compatibility. Sequences of primers and probes and its conditions are shown in **Table S1**.

### Thermal cycling conditions used for UltraPCR

All thermal cycling is performed on Bio-Rad C1000 thermal cycler with a deep well configuration with a ramp rate set at 2 °C/s. PCR thermal amplification condition is 95 °C for 2 min, followed by 40 cycles of 95 °C for 20 s and 55 - 60 °C for 30 to 60 s and 95 °C for 30 s and finally held at 12 °C.

### Fluorophore characterization

Ten fluorophores were tested in our experiments, including 3 pairs of fluorophores with similar excitation/emission spectra (FAM vs Alexa 488, Bodipy TMR-X vs HEX, Cy5 vs Atto 647N), 2 large stoke shift fluorophores (Dy-485XL, Dy-520XL), TAMRA, and Atto590. TaqMan probes conjugated with these 10 fluorophores were used in UltraPCR experiments for fluorophore characterization.

Photobleaching scans were performed on FAM vs. Alexa 488, Bodipy TMR-X vs. HEX, and Cy5 vs. Atto 647N samples in corresponding channels to confirm that FAM, Bodipy TMR-X, and Cy5 were photobleachable, while Alexa 488, HEX, and Atto 647N were photostable (defined as minimal decrease in fluorescence intensity after up to 7 repeated scans).

### 10-color TaqMan multiplexing

To show compatibility of 10 different fluorescent dyes in our 4-color system, A 10-plex panel was designed to include targets from the human mRNA transcripts (CCL3L1, PLGA1, ATF4, GAPDH, CD7), N1 region from SARS-CoV-2, genomic loci on human chromosomes 13 and 18 used in Shum et al (19), synthetic template with probe binding site (not mappable to any model organism genomes), and gene prfA from *L. cytomonogenes*. Each target was labeled with 1 of the 10 fluorescent dyes conjugated to hydrolysis probes. Each stock tube of synthetic template of each target was quantified by UltraPCR to obtain copies/uL. Next, all 10 synthetic templates, forward primers, reverse primers, and probes were assayed in one tube in duplicates (10-plex). Separately, individual synthetic templates with their respective forward primers, reverse primers, and probes were assayed in duplicate (1-plex). After emulsion generation and thermal cycling, image analysis was performed where positive partitions were counted and characterized by fluorescence level in 10 channels. The dimensional reduction software package UMAP was used to reduce the 10 channels (10D) to clusters along a 2D axis. The clusters were used to generate counts and 1-plex counts were compared to their respective clusters in the 10-plex samples.

### Comboplex studies

For comboplex with hydrolysis probes, probes targeting the same sequence with different fluorescent dyes were used so that each target could be labeled with multiple colors. Synthetic template for prfA, primers and 1 to 3 hydrolysis probes were added in the mixture and followed hydrolysis probe amplifying condition.

For the 22-plex comboplex experiments, UltraPCR’s Universal Multiplex (UM) Kit and UltraPCR Mix (Enumerix) were used to perform multiplexing without the use of hydrolysis probes. UM technology utilizes a 5’ adapter to a PCR forward primer to label targets. Targets were labeled with either 1 color or 2 colors using different combinations of forward primers. For 1-color labeling, 1 forward primer per target was used and for 2-color labeling, 2 forward primers per target were used. Once minimal nonspecific interactions between all primers and probes were confirmed without template addition, all the primers and probes for the 22 targets were pooled together respectively. Final primer concentration for targets labeled with 2 colors were 20 nM each for the two forward primers and 200 nM for the reverse primer. For targets labeled with 1 color, the final concentrations were 40 nM and 200 nM for the forward and reverse primer, respectively. Each synthetic target template was added accordingly depending on 4 plex, 8 plex and 22 plex conditions. Annealing temperature in UM thermal cycling condition was increased from 56 °C to 60 °C to minimize non-specific amplifications.

## Author Contributions

J.H.L., J.W.K., M.M., S.L., S.P.A.F., H.C.F. and E.Y.S conceptualized, designed, and developed the UltraPCR and comboplex technology and research. H.G.L., E.J.B., J.S., V.K.R.. performed the research. E.Y.S., J.H.L., M.M., H.G.L., J.S., C.L.B analyzed the data. E.Y.S. and H.C.F. wrote the manuscript.

## Notes

Competing Interest Statement: The authors declare the following competing financial interest(s): J.H.L., J.W.K., H.G.L., M.M., E.J.B.,S.L.,J.S., V.K.R, C.L.B., S.P.A.F., H.C.F. and E.Y.S. are employees of Enumerix, Inc., a company that commercializes DNA counting technologies. This work was supported by Enumerix, Inc. and National Institute of General Medical Sciences of the National Institue of Health under the grant number 1R44GM149104-01. In addition, E.Y.S., J.H.L., S.L., S.P.A.F., H.C.F., J.W.K., H.G.L. are inventors on patents and patent applications relating to methods described in this manuscript.

## ACKNOWLEDGMENT

We thank Mr. Ari Chaney, Ms. Kat George, Mr. Aaron Solomon, and Mr. Ivan Wong for their support of this research.

## REFERENCES

1. P. W. Chiang, et al., Use of a fluorescent-PCR reaction to detect genomic sequence copy number and transcriptional abundance. Genome Res. 6, 1013–1026 (1996).

2. M. Adinolfi, J. Sherlock, B. Pertl, Rapid detection of selected aneuploidies by quantitative fluorescent PCR. BioEssays 17, 661–664 (1995).

3. M. Adinolfi, B. Pertl, J. Sherlock, Rapid detection of aneuploidies by microsatellite and the quantitative fluorescent polymerase chain reaction. Prenat. Diagn. 17, 1299–1311 (1997).

4. I. Findlay, P. Quirke, J. Hall, A. Rutherford, Fluorescent PCR: A new technique for PGD of sex and single-gene defects. J. Assist. Reprod. Genet. 13, 96–103 (1996).

5. K. Mullis, et al., Specific Enzymatic Amplification of DNA In Vitro: The Polymerase Chain Reaction. Cold Spring Harb. Symp. Quant. Biol. 51, 263–273 (1986).

6. M. C. Edwards, R. A. Gibbs, Multiplex PCR: advantages, development, and applications. Genome Res. 3, S65–S75 (1994).

7. I. Saito, B. Servenius, T. Compton, R. I. Fox, Detection of Epstein-Barr virus DNA by polymerase chain reaction in blood and tissue biopsies from patients with Sjogren’s syndrome. J. Exp. Med. 169, 2191–2198 (1989).

8. R. K. Saiki, et al., Primer-Directed Enzymatic Amplification of DNA with a Thermostable DNA Polymerase. Science 239, 487–491 (1988).

9. R. K. Saiki, et al., Enzymatic Amplification of β-Globin Genomic Sequences and Restriction Site Analysis for Diagnosis of Sickle Cell Anemia. Science 230, 1350–1354 (1985).

10. J. S. Chamberlain, R. A. Gibbs, J. E. Rainer, P. N. Nguyen, C. Thomas, Deletion screening of the Duchenne muscular dystrophy locus via multiplex DNA amplification. Nucleic Acids Res. 16, 11141–11156 (1988).

11. E. Zietkiewicz, M. Labuda, D. Sinnett, F. H. Glorieux, D. Labuda, Linkage mapping by simultaneous screening of multiple polymorphic loci using Alu oligonucleotide-directed PCR. Proc. Natl. Acad. Sci. 89, 8448–8451 (1992).

12. A. Ballabio, JoelE. Ranier, JeffreyS. Chamberlain, M. Zollo, C. T. Caskey, Screening for steroid sulfatase (STS) gene deletions by multiplex DNA amplification. Hum. Genet. 84 (1990).

13. P. Markoulatos, N. Siafakas, M. Moncany, Multiplex polymerase chain reaction: A practical approach. J. Clin. Lab. Anal. 16, 47–51 (2002).

14. TaqMan vs SYBR Chemistry - US. (October 4, 2023). ThermoFisher Scientific. https://www.thermofisher.com/us/en/home/life-science/pcr/real-time-pcr/real-time-pcr-learning-center/real-time-pcr-basics/taqman-vs-sybr-chemistry-real-time-pcr.html

15. M. Arya, et al., Basic principles of real-time quantitative PCR. Expert Rev. Mol. Diagn. 5, 209–219 (2005).

16. M. Buh Gašparič, et al., Comparison of nine different real-time PCR chemistries for qualitative and quantitative applications in GMO detection. Anal. Bioanal. Chem. 396, 2023–2029 (2010).

17. E. Navarro, G. Serrano-Heras, M. J. Castaño, J. Solera, Real-time PCR detection chemistry. Clin. Chim. Acta 439, 231–250 (2015).

18. Y. N. Teo, E. T. Kool, DNA-Multichromophore Systems. Chem. Rev. 112, 4221–4245 (2012).

19. E. Y. Shum, et al., Next-Generation Digital Polymerase Chain Reaction: High-Dynamic-Range Single-Molecule DNA Counting via Ultrapartitioning. Anal. Chem. 94, 17868–17876 (2022).

20. B. J. Hindson, et al., High-Throughput Droplet Digital PCR System for Absolute Quantitation of DNA Copy Number. Anal. Chem. 83, 8604–8610 (2011).

21. A. Tiwari, et al., Application of digital PCR for public health-related water quality monitoring. Sci. Total Environ. 837, 155663 (2022).

22. A. S. Basu, Digital assays part I: partitioning statistics and digital PCR. SLAS Technol. 22, 369–386 (2017).

23. P. Liao, et al., Three-dimensional digital PCR through light-sheet imaging of optically cleared emulsion. Proc. Natl. Acad. Sci. 117, 25628–25633 (2020).

24. QPCR Guidelines: Multiplex Dye Compatibility. (n.d). Agilent. https://www.agilent.com/files/Mobio/QPCR%20Guidelines_Multiplex_Dye_Compatibility.pdf

25. A. S. Whale, J. F. Huggett, S. Tzonev, Fundamentals of multiplexing with digital PCR. Biomol. Detect. Quantif. 10, 15–23 (2016).

26. J. F. Huggett, et al., The Digital MIQE Guidelines: Minimum Information for Publication of Quantitative Digital PCR Experiments. Clin. Chem. 59, 892–902 (2013).

27. M. Ester, H.-P. Kriegel, J. Sander, X. Xu, A Density-Based Algorithm for Discovering Clusters in Large Spatial Databases with Noise. kdd 96, 226–231.

28. C. Kempf, et al., Tissue Multicolor STED Nanoscopy of Presynaptic Proteins in the Calyx of Held. PLoS ONE 8, e62893 (2013).

29. B. Zhang, C. Kang, D. R. Davydov, Conformational Rearrangements in the Redox Cycling of NADPH-Cytochrome P450 Reductase from Sorghum bicolor Explored with FRET and Pressure-Perturbation Spectroscopy. Biology 11, 510 (2022).

30. T. H. Linz, W. Hampton Henley, J. Michael Ramsey, Photobleaching kinetics-based bead encoding for multiplexed bioassays. Lab. Chip 17, 1076–1082 (2017).

31. F. Schlenker, et al., Virtual Fluorescence Color Channels by Selective Photobleaching in Digital PCR Applied to the Quantification of KRAS Point Mutations. Anal. Chem. 93, 10538–10545 (2021).

32. L. McInnes, J. Healy, J. Melville, UMAP: Uniform Manifold Approximation and Projection for Dimension Reduction (2018) 10.48550/ARXIV.1802.03426 (June 30, 2023).

33. S. H. Lee, et al., Performance of a multiplex PCR pneumonia panel for the identification of respiratory pathogens and the main determinants of resistance from the lower respiratory tract specimens of adult patients in intensive care units. J. Microbiol. Immunol. Infect. 52, 920–928 (2019).

34. J. Madic, et al., Three-color crystal digital PCR. Biomol. Detect. Quantif. 10, 34–46 (2016).

35. P.-L. Quan, M. Sauzade, E. Brouzes, dPCR: A Technology Review. Sensors 18 (2018).

36. B. Vogelstein, K. W. Kinzler, Digital PCR. Proc. Natl. Acad. Sci. 96, 9236–9241 (1999).

37. G. Pohl, I.-M. Shih, Principle and applications of digital PCR. Expert Rev. Mol. Diagn. 4, 41–47 (2004).

38. J. Madic, C. Jovelet, I. Dehri, A. C. Mallory, “6-Color Crystal Digital PCRTM for the High-Plex Detection of EGFR Mutations in Non-Small Cell Lung Cancer” in Lung Cancer, P. G. Santiago-Cardona, Ed. (Springer US, 2021), pp. 127–144.

39. FACSymphony™ A5 | High-Parameter Cell Analyzer (October 5, 2023). BD Biosciences. https://www.bdbiosciences.com/en-us/products/instr uments/flow-cytometers/research-cell-analyzers/bd-f acsymphony-a5

40. R. M. Ferrie, et al., Development, multiplexing, and application of ARMS tests for common mutations in the CFTR gene. Am. J. Hum. Genet. 51, 251–262 (1992).

41. D. Sint, L. Raso, M. Traugott, Advances in multiplex PCR: balancing primer efficiencies and improving detection success. Methods Ecol. Evol. 3, 898–905 (2012).

42. B. Leatham, et al., A rapid, multiplex digital PCR assay for EGFR, KRAS, BRAF, ERBB2 variants and ALK, RET, ROS1, NTRK1 gene fusions in non-small cell lung cancer (2023) 10.1101/2023.03.09.531949 (June 27, 2023).

43. A. Rajagopal, et al., Significant Expansion of Real-Time PCR Multiplexing with Traditional Chemistries using Amplitude Modulation. Sci. Rep. 9, 1053 (2019).

44. L. Jacky, et al., Virtual-Partition Digital PCR for High-Precision Chromosomal Counting Applications. Anal Chem 93, 17020–17029 (2021).

45. L. Jacky, et al., Robust Multichannel Encoding for Highly Multiplexed Quantitative PCR. Anal Chem 93, 4208–4216 (2021).

46. S. Dube, J. Qin, R. Ramakrishnan, Mathematical Analysis of Copy Number Variation in a DNA Sample Using Digital PCR on a Nanofluidic Device. PLoS ONE 3, e2876 (2008).

47. S. R. Kim, C.-S. Ki, N. Y. Lee, Rapid detection and identification of 12 respiratory viruses using a dual priming oligonucleotide system-based multiplex PCR assay. J. Virol. Methods 156, 111–116 (2009).

48. A. A. El Kholy, et al., The use of multiplex PCR for the diagnosis of viral severe acute respiratory infection in children: a high rate of co-detection during the winter season. Eur. J. Clin. Microbiol. Infect. Dis. 35, 1607–1613 (2016).

49. A. Stahlberg, L. Stein, T. E. Godfrey, Simple, multiplexed, PCR-based barcoding of DNA enables sensitive mutation detection in liquid biopsies using sequencing. Nucleic Acids Res. 44 (2016).

50. P. Song, et al., Limitations and opportunities of technologies for the analysis of cell-free DNA in cancer diagnostics. Nat. Biomed. Eng. 6, 232–245 (2022).

51. T. Koressaar, M. Remm, Enhancements and modifications of primer design program Primer3. Bioinformatics 23, 1289–1291 (2007).

52. T. Kõressaar, et al., Primer3_masker: integrating masking of template sequence with primer design software. Bioinformatics 34, 1937–1938 (2018).

53. A. Untergasser, et al., Primer3—new capabilities and interfaces. Nucleic Acids Res. 40, e115–e115 (2012).

