## Supplementary for "New realm of precision multiplexing enabled by massively-parallel single molecule UltraPCR"

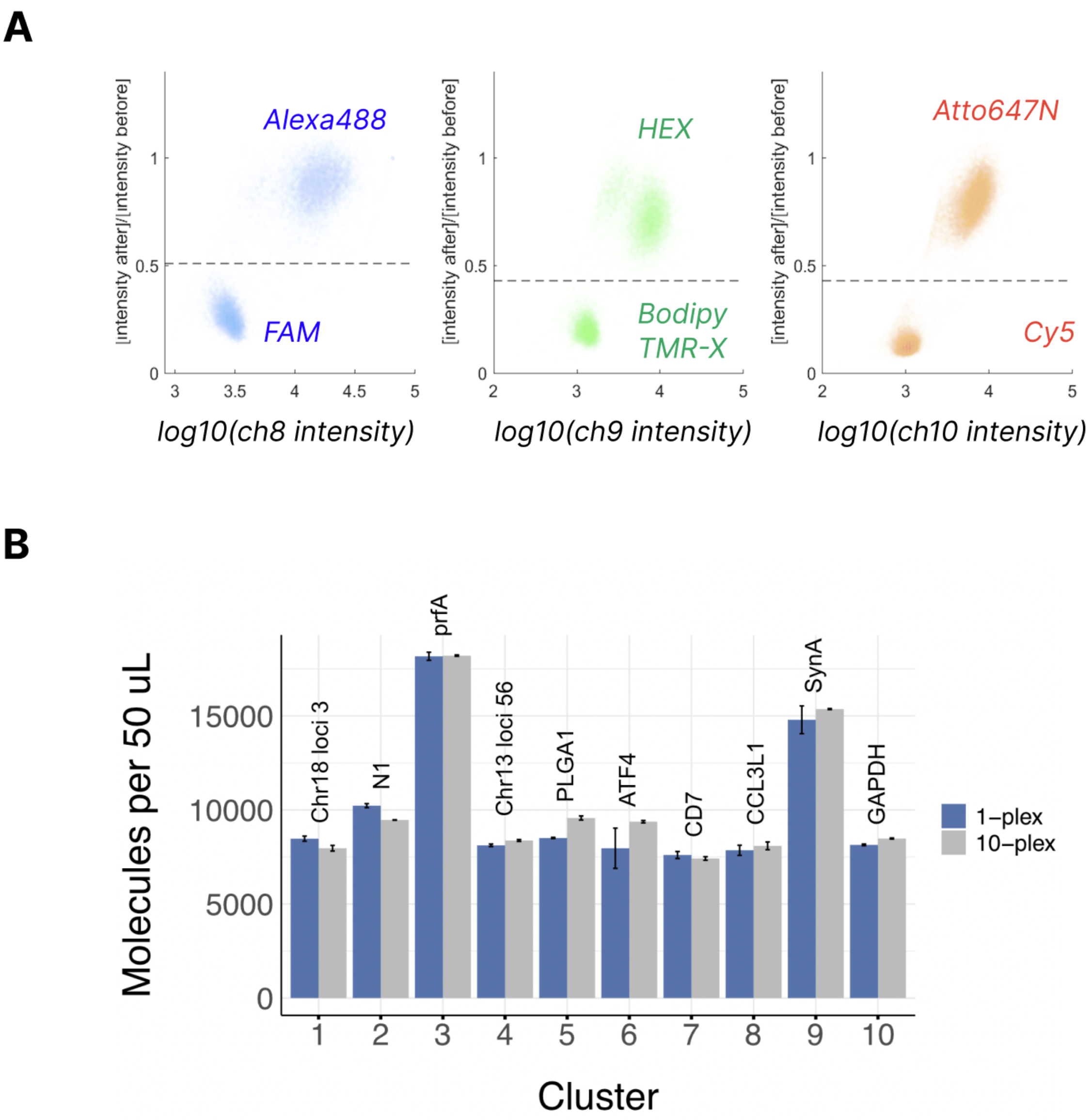


**Figure S1. Expanding dye repertoire in UltraPCR and achieving a 10-dye multiplex assay.** (A) Scatterplot of partitions where x-axis is the fluorescence intensity measuring the corresponding channel (post photobleaching), and y-axis represents the ratio of fluorescence intensities before and after photobleaching. Partitions with dyes that are photobleachable appear closer to the bottom left of the graph and can be automatically gated away from nonphotobleachable partitions. (B) Comparison in UltraPCR counting accuracy for each target when each is amplified and counted alone (1-plex) or when multiplexed together and counted using UMAP/clustering as shown in Figure 2. Error bar denotes standard error (n=2).

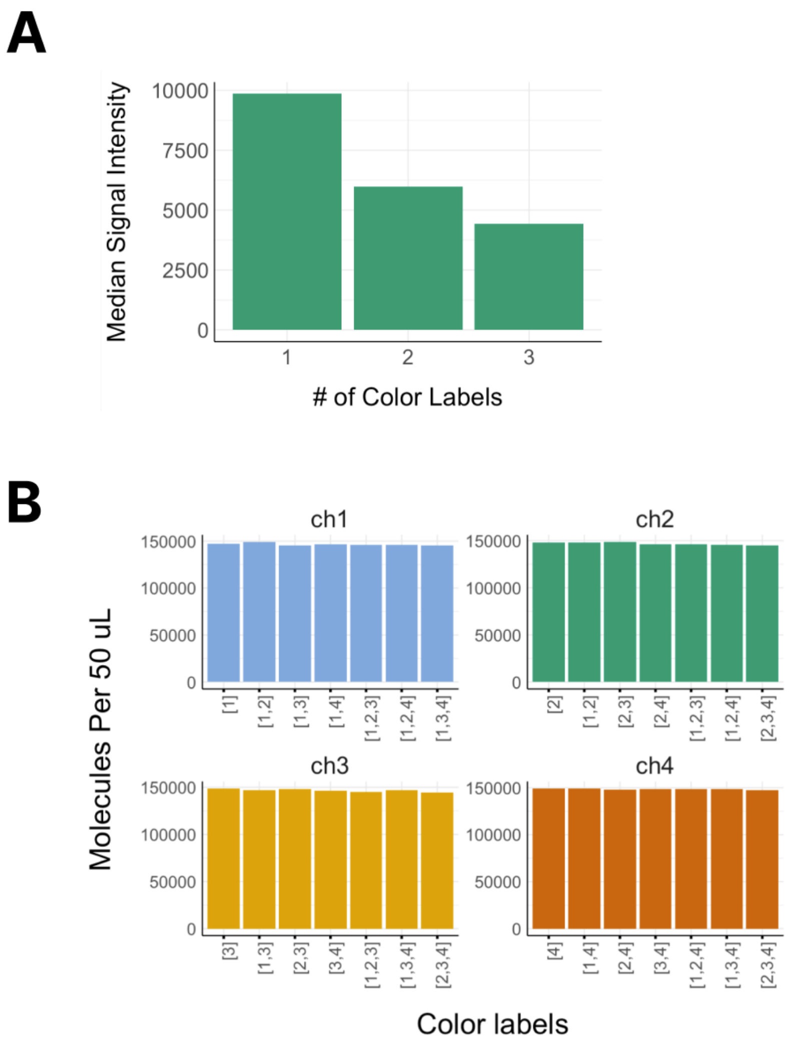


**Figure S2. Proof of concept studies for comboplex.** (A) Fluorescence signaling ch2, as measured by the median signal intensity of positive partitions, for *prfA*, showing that in some assay designs, the increase in *k* can reduce intensity. However, since UltraPCR occurs in singlet realm, it does not affect the auto thresholding section of the UltraPCR Imager analysis pipeline. (B) Molecule counts for each comboplex configuration to measure *prfA*, showcasing that precision counting is preserved whether $k$ = 1, 2, or 3 in UltraPCR.

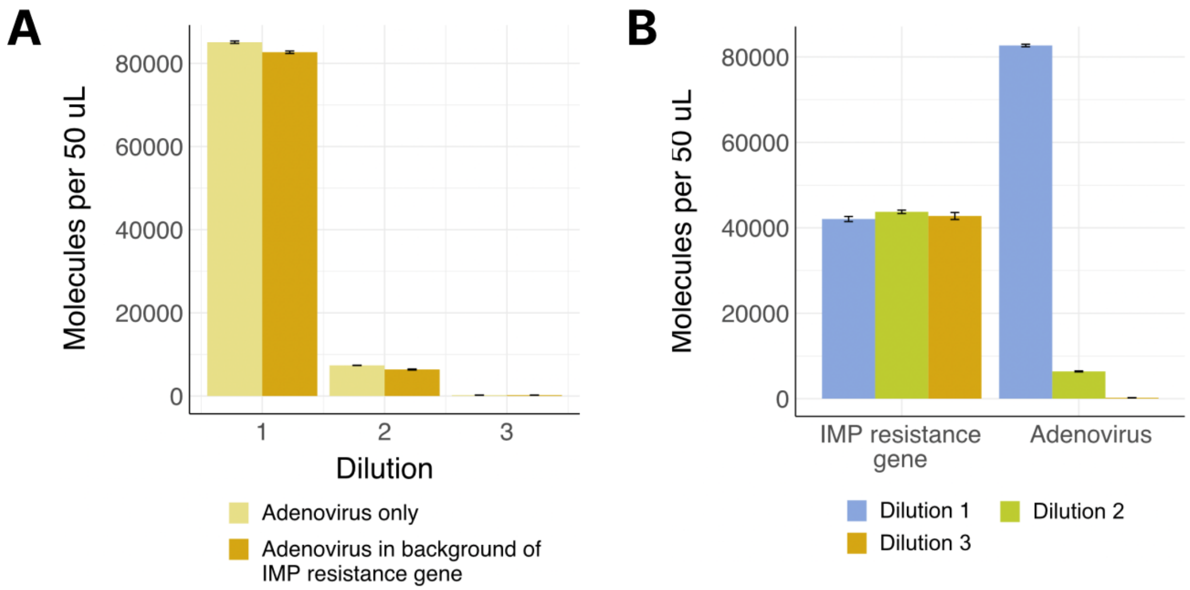


**Figure S3. Titration studies using the 22-plex comboplex pathogen ID panel**. Bar graphs showing the number of molecules detected for spiked ADV54 (Adenovirus) DNA in the co-presence of an antibiotic resistance gene Imp. In the first configuration of data display shown in (A) serially diluted ADV54 counts remain the same with or without Imp DNA spiked in. (B) In the same serial dilution experiment of ADV54, the level of Imp gene remains unchanged. Together it shows how simultaneous detection of targets – though in the same assay – is counted independently from each other regardless of concentration of each target.

**Table S1. Probes and gene specific primers sequences used in the paper**

| Name | Sequence |
|---|---|
| N1-FAM | ACCCCGCATTACGTTTGGTGGACC |
| Chr13-Alexa488 | C+C+CA+C+CTC+CA |
| CCL3L-HEX | TTCGAGGCCCAGCGACCTCA |
| Chr18-Atto590 | C+T+C+CT+CA+G+CA |
| SynA-Cy5 | CAAGCAGAAGACGGCATACGAGAT |
| PLGA1-Dy485XL | TGGATACAGAGGGCCAACTGTATTTAGGA |
| ATF4-Dy520XL | TGCCTTCTCCGGGACAGATTGGAT |
| GAPDH-Bodipy TMR-X | CCCATCACCATCTTCCAGGAGCGA |
| CD7-TAMRA | ACTGTGCTCGTGGCGGGATAAGAA |
| prfA-FAM/HEX/TEX615 | AGCCAACCGATGTTTCTGTATCAA |
| N1-F | GACCCCAAAATCAGCGAAAT |
| N1-R | TCTGGTTACTGCCAGTTGAATCTG |
| Chr13-1-F | GCCTGGTGATGACCTCTGTC |
| Chr13-1-R | CTGGTTGTCACTCACTCCCC |
| CCL3L1-F | GGGTCCAGAAATACGTCAGT |
| CCL3L1-R | CATGTTCCCAAGGCTCAG |
| Chr18-1-F | ACTCCAGCCTTTAGAGAAAAATGC |
| Chr18-1-R | TGAGAGTGCCACCTACAAACA |
| MPV-F | CAGCAAGCTCACCAGAGACA |
| MPV-R | GTGTTGTGTGTGTGTCGACG |
| PLGA1-F | GGTTGAATTTGTGGATGTGGAA |
| PLGA1-R | TCTGCTTATTCACATATTCTGGTTTGA |
| ATF4-F | GTCAGTCCCTCCAACAACAGC |
| ATF4-R | GTCATCTATACCCAACAGGGC |
| GAPDH-F | GTCAAGGCTGAGAACGGGAA |
| GAPDH-R | AAATGAGCCCCAGCCTTCTC |
| CD7-F | TGCTGGCGAGGACACAGATAA |
| CD7-R | TGCGACATGTCCTCGTACAC |
| prfA-F | CCGCAAATAGAGCCAAGCTT |
| prfA-R | GTGAGAACGGGACCATCATG |
| Enterobacter_F | GCGGTAATCAAAGTCGGTGC |
| Paraflu4b__F | AGAGCAAGCTCCCGTAATCG |
| Influenza_A virus_F | CAGCATCGGTCTCACAGACA |
| Escherichia coli_F | AACTGTATACCGCACCTGGC |
| Rhino_F | CACAAGGACCAATAGCCGGT |

| | |
|---|---|
| Paraflu1_F | GATCCTCAGGTTGGCACTCC |
| Paraflu2_F | AGATGTGTGGCCGCTTAACA |
| Bordetella_F | CGAGAAAAGCGGATCGAGGA |
| Rsv_F | TTGGATCTGCAATCGCCAGT |
| Chlamydia_F | CATTTGCTGGTTCTGTCGGC |
| Coro229E_F | GGCAAACGGGTGGATTTGTC |
| Adv54_F | AAGTCAGCAACTACCCCGTG |
| Mpv_F | CAGCAAGCTCACCAGAGACA |
| Rhino_F | CACAAGGACCAATAGCCGGT |
| Paraflu1_F | GATCCTCAGGTTGGCACTCC |
| Paraflu2_F | AGATGTGTGGCCGCTTAACA |
| Bordetella_F | CGAGAAAAGCGGATCGAGGA |
| Rsv_F_F | TTGGATCTGCAATCGCCAGT |
| Chlamydia_F | CATTTGCTGGTTCTGTCGGC |
| Coro229E_F | GGCAAACGGGTGGATTTGTC |
| Adv54_F | AAGTCAGCAACTACCCCGTG |
| Mpv_F | CAGCAAGCTCACCAGAGACA |
| Legionella pneumophila_F | TGACTGCAGCTGTTATGGGG |
| Klebsiella oxytoca_F | CATCGAAGCCAGCACATTCG |
| Staphylococcus aureus_F | TGCAGCTCAACATATCACACCT |
| Serratia marcescens_F | TGCTGAAAGTGACCGACCAG |
| Proteus mirabili_F | ACCGAAGAGCGTTGGTCAAT |
| Klebsiella pneumoniae.2_F | CAACCAGTTTGCCAGACACG |
| IMP_F | TGGGGCGTTGTTCCTAAACA |
| KPC_F | TTCCCACTGTGCAGCTCATT |
| NDM_F | TTGCGACTTATGCCAATGCG |
| Legionella pneumophila_F | TGACTGCAGCTGTTATGGGG |
| Klebsiella oxytoca_F | CATCGAAGCCAGCACATTCG |
| Staphylococcus aureus_F | TGCAGCTCAACATATCACACCT |
| Serratia marcescens_F | TGCTGAAAGTGACCGACCAG |
| Proteus mirabili_F | ACCGAAGAGCGTTGGTCAAT |
| Klebsiella pneumoniae.2_F | CAACCAGTTTGCCAGACACG |
| IMP_F | TGGGGCGTTGTTCCTAAACA |
| KPC_F | TTCCCACTGTGCAGCTCATT |
| NDM_F | TTGCGACTTATGCCAATGCG |
| Enterobacter_R | CAGCTACCACGCCTTCTTCA |
| Paraflu4b_R | GGGTCTCGCTAACGGATCAA |

| | |
|---|---|
| Influenza_A virus_R | TGTTCCATAGCCTTTGCCGT |
| Escherichia coli_R | AACGGCCACCAGAGTTACTG |
| Rhino_R | TCGCTGTTGTTTTTGCCCTG |
| Paraflu1_R | TTGCAGTCTGGGTTTCCTGG |
| Paraflu2_R | GGGATTGGTTCGGTTGGACT |
| Bordetella_R | CGCTGTACCCATCTCCAGAC |
| Rsv_R | ACTACGGCCTTGTTTGTGGA |
| Chlamydia_R | GCAGCACCCTCCCATATTGT |
| Coro229E_R | CCCAGACGACACCTTCAACA |
| Adv54_R | GTAGACGGCGAGATCGTTGT |
| Mpv_R | GTGTTGTGTGTGTGTCGACG |
| Legionella pneumophila_R | TCTTTCCCAAATCGGCACCA |
| Klebsiella oxytoca_R | GCGTGACGGTACAGCTATCA |
| Staphylococcus aureus_R | CGGAATCTGATGTTGCAGTTGT |
| Serratia marcescens_R | CCTCTTCCAGCGAGTGCATA |
| Proteus mirabili_R | ACCGGTTTCAAGCAGTTCTT |
| Klebsiella pneumoniae.2_R | GGGACATGAGTCGACCGAAA |
| IMP_R | CACGCTCCACAAACCAAGTG |
| KPC_R | CCACAGAACCAGCGCATTTT |
| NDM_R | GAAAGTCAGGCTGTGTTGCG |

LNA nucleotides are marked with plus before the corresponding base.
